# Why do scientists fabricate and falsify data? A matched-control analysis of papers containing problematic image duplications

**DOI:** 10.1101/126805

**Authors:** Daniele Fanelli, Rodrigo Costas, Ferric C. Fang, Arturo Casadevall, Elisabeth M. Bik

## Abstract

It is commonly hypothesized that scientists are more likely to engage in data falsification and fabrication when they are subject to pressures to publish, when they are not restrained by forms of social control, when they work in countries lacking policies to tackle scientific misconduct, and when they are male. Evidence to test these hypotheses, however, is inconclusive due to the difficulties of obtaining unbiased data.

Here we report a pre-registered test of these four hypotheses, conducted on papers that were identified in a previous study as containing problematic image duplications through a systematic screening of the journal PLoS ONE. Image duplications were classified into three categories based on their complexity, with category 1 being most likely to reflect unintentional error and category 3 being most likely to reflect intentional fabrication. Multiple parameters connected to the hypotheses above were tested with a matched-control paradigm, by collecting two controls for each paper containing duplications.

Category 1 duplications were mostly not associated with any of the parameters tested, in accordance with the assumption that these duplications were mostly not due to misconduct. Category 2 and 3, however, exhibited numerous statistically significant associations. Results of univariable and multivariable analyses support the hypotheses that academic culture, peer control, cash-based publication incentives and national misconduct policies might affect scientific integrity. Significant correlations between the risk of image duplication and individual publication rates or gender, however, were only observed in secondary and exploratory analyses.

Country-level parameters generally exhibited effects of larger magnitude than individual-level parameters, because a subset of countries was significantly more likely to produce problematic image duplications. Promoting good research practices in all countries should be a priority for the international research integrity agenda.

## INTRODUCTION

The scientific literature is plagued by a small yet not negligible percentage of papers with fabricated or falsified results. Survey studies suggest that 1-2% of scientists admit to having consciously fabricated or falsified data at least once [1, 2], although the actual percentage of fabricated papers might be just a fraction of the percentage of self-reported misconduct, at least in the field of Psychology [3]. Direct assessments of the rate of image manipulation in biology, however, suggest that between 1-4% of papers contain problematic images, at least part of which is likely to result from intentional fabrication [4, 5].

Multiple sociological, cultural and psychological factors are hypothesized to increase the risk that scientists engage in scientific misconduct, and testing these hypotheses is a matter of ongoing theoretical and empirical research. Particular attention has been paid to four major factors:

- *Pressures to publish:* it is commonly suggested that scientists might engage in scientific misconduct in response to high expectations of productivity and/or impact. This concern has already guided numerous policies and initiatives aimed at discouraging scientists from publishing too much and/or from pursuing high impact at all costs (e.g. [6-8]). Pressures to publish may be higher and increasing in countries in which institutions are evaluated based on their publication performance (e.g. United Kingdom’s Research Excellence Framework), and/or in countries in which career advancement is determined by publications (e.g. tenure-track system in the United States of America) and/or in countries in which high-profile researchers are rewarded with cash (e.g. reward policies in China, see [9]). The pressures to publish hypothesis is supported by perceptions reported in anonymous surveys [10, 11], but fails to predict the incidence of retractions and corrections [12], historical trends of scientists’ publication rate [13], and the likelihood to report over-estimated effects [14].
- *Social control:* sociological and psychological theories suggest that individuals are less likely to engage in misconduct when scrutiny of their work is ensured by peers, mentors or society (e.g. [15]). An elaborate socio-economic hypothesis predicts that mutual criticism and policing of misconduct might be least likely to occur in developing countries in which academia was built on the German model, and might be most likely in developed (i.e. highly regulated) countries with an Anglo-American (i.e. highly egalitarian) academic culture [16]. Within teams, the social control hypothesis predicts that mutual criticism is likely to be directly proportional to the number of team members and inversely to their geographic distance, a prediction supported by studies on retractions and bias in the literature [12, 17].
- *Misconduct policies:* a growing number of countries and/or institutions are establishing official policies that define scientific misconduct and that regulate how suspected cases can be identified, investigated and punished. These policies express the rationale that clear rules and sanctions will have a deterrent effect on misconduct [18]. Countries differ widely in how they define and enforce misconduct policies, and it is commonly suggested that the greatest deterrent effect would be obtained by misconduct policies that are legally enforced e.g. [19].
- *Gender:* males are more prone to taking risk and more status-oriented than females, and might therefore be more likely to engage in scientific misconduct [20]. This hypothesis received some support by statistics about findings of misconduct by the US Office of Research Integrity [20]. However, other interpretations of these data have been proposed [21] and gender did not significantly predict the likelihood to produce a retracted or corrected paper, once various confounders were adjusted for in a matched-control analysis [12].

Progress in assessing the validity of these hypotheses in explaining the prevalence of misconduct has been hampered by difficulties in obtaining reliable data. A primary source of evidence about scientific misconduct is represented by anonymous surveys. These however are very sensitive to methodological choices and, by definition, report what a sample of voluntary respondents think and are willing to declare in surveys –not necessarily what the average scientist actually thinks and does [1, 3, 22]. Retractions of scientific papers, most of which are due to scientific misconduct [23], offer a pool of actual cases whose analyses have given important insights (e.g. [12, 23-25]). Results obtained on retractions, however, may not be generalizable, because retractions still constitute a very small fraction of the literature and by definition are the result of a complex process that can be influenced by multiple contingent factors, such as level of scrutiny of a literature, presence of retraction policies in journals, and the scientific community’s willingness to act [26].

An unprecedented opportunity to probe further into the nature of scientific misconduct is offered by a recent dataset of papers that contain image duplications of a questionable or manifestly fraudulent nature, i.e. Bik et al. 2016 [5]. These papers were identified by direct visual inspection of 20,621 papers that contained images of Western Blots, nucleic acid gels, flow cytometry plots, histopathology or other forms of image that were published between the years 1995 and 2015 in 40 journals. Having been obtained by a systematic screening of the literature, this sample is free from most limitations and biases that affect survey and retraction data, and therefore offers a representative picture of errors and/or misconduct in the literature – at least with regard to image duplications in biological research. Descriptive analyses of these data have yielded new insights into the rate of scientific misconduct and its relative prevalence amongst different countries.

We conducted a pre-registered analysis (osf.io/w53yu) of data from [5] to test, using a matched-control approach, multiple postulated social and psychological risk factors for scientific misconduct. Our analysis focused on the largest and most homogeneous subsample of the original data set, i.e. N=346 papers with duplicated images identified from a sample of 8,138 papers published in the journal *PLoS ONE*, between the years 2013 and 2014.

Image duplications included in our sample could be due to unintentional error, questionable practice or outright scientific misconduct. Following the classification used in [5], image duplications were grouped in three categories according to their complexity and therefore their likelihood to result from scientific misconduct:

- *Category 1:* Simple duplications, in which the same image is presented twice to represent different conditions, possibly due to accidental mislabeling (N=83).
- *Category 2:* Duplication with re-positioning, in which one image has been shifted, rotated or reversed, suggesting some level of active intervention by the researcher (N=186)
- *Category 3:* Duplication with alteration, in which figures contained evidence of cutting, patching and other forms of substantive embellishment and manipulation which betrays a possible intention to mislead (N=77).

Category 1 duplications are most likely to due to error, whilst categories 2 and 3 are likely to contain a mixture of errors and intentional fabrications. Therefore, if factors predicted to affect scientific misconduct have any effect at all, such effects are predicted to be most relevant in category 2 and 3 duplications and to have little or no effect on category 1 errors.

For each paper containing duplicated images, we identified two controls that had been published in the same journal and time period, and that contained images of Western blots without detectable signs of duplication. We then measured a set of variables that were relevant to each of the hypotheses listed above, and used logistic regression to test whether and how these variables were associated with the risk of committing scientific misconduct.

## RESULTS

Figure 1 reports the effects in each category of duplication of each tested parameter (i.e. odds ratio and 95% confidence interval), grouped by each composite hypothesis, with an indication of the direction of effect predicted by that hypothesis. In line with our overall predictions, Category 1 duplications yielded a null association with nearly all of the parameters tested (Figure 1, green error bars), and/or yielded markedly different effects from Category 2 and Category 3 papers (Figure 1, orange and red bars, respectively). Sharp and highly significant differences between effects measured on the latter and the former duplication categories were observed for authors’ citation scores and journal scores (Fig 1a), and for several country-level and team-level parameters (i.e. Fig 1 b-e). No significant difference was observed amongst gender effects, except for a tendency of Category 3 duplications to be more common amongst female authors (Fig 1f).

**Fig 1:**
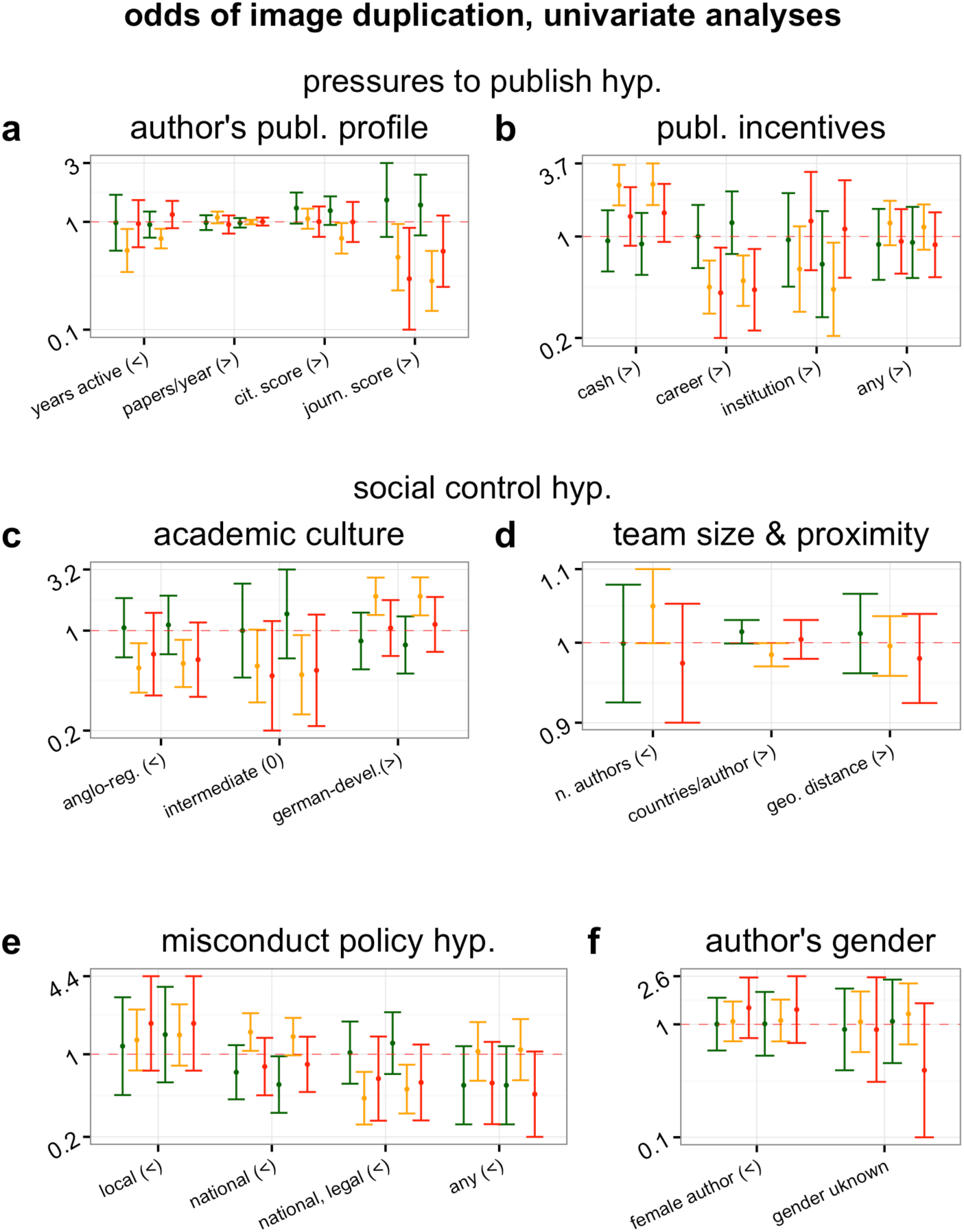
Effect (odds ratio and 95%CI) of characteristics of study and of first and last authors on the likelihood to publish a paper containing a Category 1 (green), Category 2 (yellow) or Category 3 (red) problematic image duplication. When six error bars are associated with one test, the first three error bars correspond to data from the first author and the last three are for data from the last author. Panels are subdivided according to overall hypothesis tested, and signs in parentheses indicate direction of expected effect (“>” : OR>1; “<” : OR<1; “0”: intermediate effect predicted).

Differences between effects measured on Category 2 and 3 duplications were not always consistent with our prediction that Category 3 duplications should exhibit the largest effect sizes. For example, the number of years of activity of the author was only significantly associated with Category 2 duplications (Fig 1a). In most cases, however, the confidence intervals of effects measured for Categories 2 and 3 were largely overlapping, suggesting that differences between Category 2 and 3 might be due to the smaller sample size (lower statistical power) achieved for the latter category. Overall, therefore, results of univariable analyses are consistent with our predictions and confirm the original assessment of the status of these categories suggested by Bik et al. (2016): Category 1 duplications are most likely to reflect genuine errors whilst Category 2 and 3 errors are most likely to reflect intentional manipulations. Hypotheses about determinants of scientific misconduct, therefore, are most directly testable on the latter two categories, which were combined in all subsequent analyses reported in the main text. A supplementary file reports all numerical results of all analyses reported in the main text as well as all robustness analyses obtained on each separate duplication category and on all categories combined (see SI).

Results of univariable tests combining Category 2 and 3 papers together are in good agreement with the social control hypothesis (Fig 2c) and partial agreement with the misconduct policy hypothesis (Fig 2e). The gender hypothesis was not supported (Fig 2f). The pressures to publish hypothesis was not or negatively supported by most analyses. In agreement with some predictions, the risk of misconduct was higher in countries in which publications are rewarded by cash incentives (Fig 2b) and was lower for researchers with a shorter publication time-span (i.e. presumably early-career researchers, Fig 2a). Contrary to predictions, however, the risk of misconduct was lower for authors with higher journal score (Fig 1a) and in countries with publication incentive policies that are career-based and institutional-based, despite the fact that the latter are those where pressures to publish are said to be highest [10].

**Fig 2:**
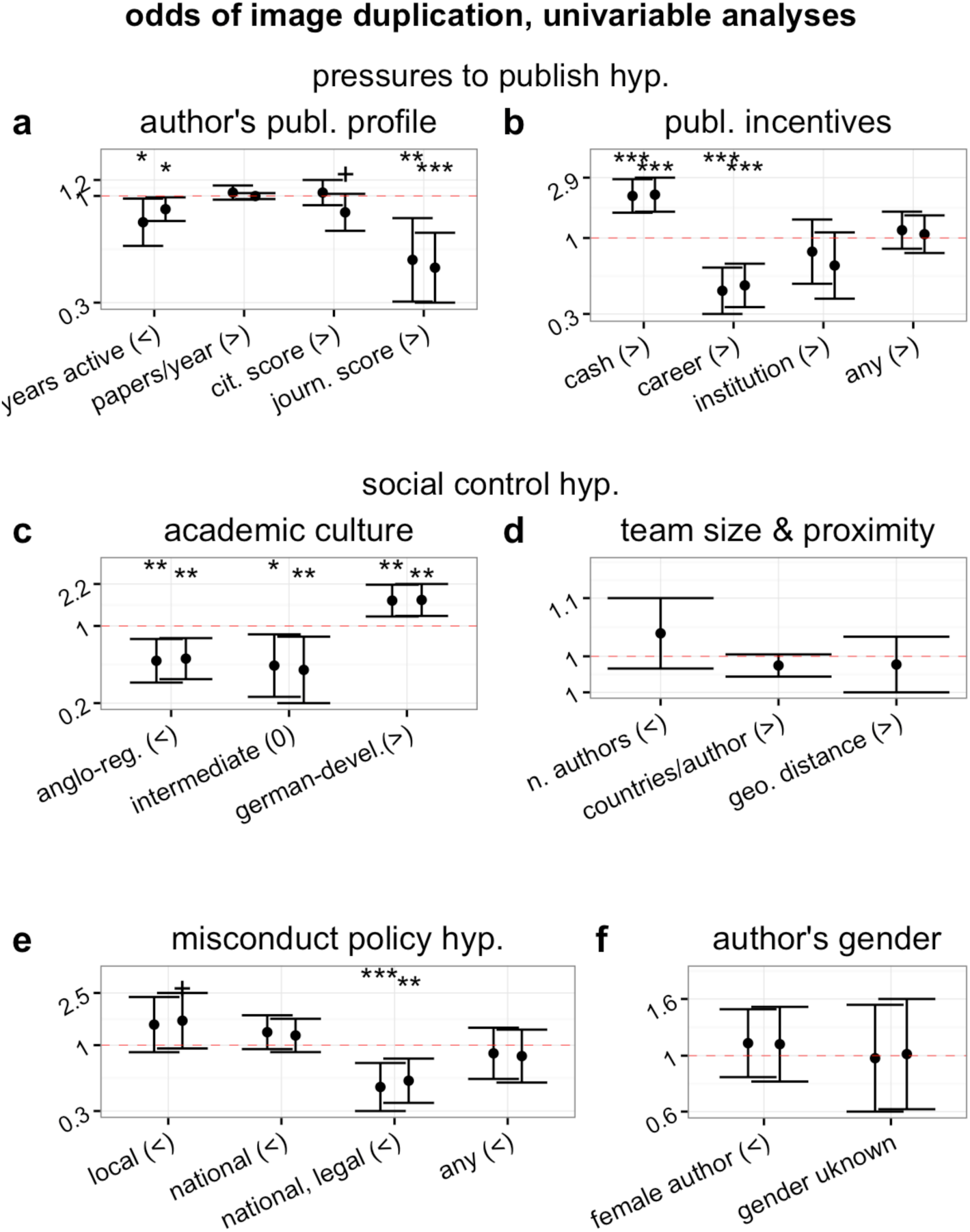
Effect (odds ratio and 95%CI) of characteristics of study and of first and last authors on the likelihood to publish a paper containing a Category 2 or 3 problematic image duplication. For each individual-level parameter, first and second error bars correspond to data from first and last authors, respectively. Panels are subdivided according to overall hypothesis tested, and signs in parentheses indicate direction of expected effect (“>” : OR>1; “<” : OR<1; “0”: intermediate effect predicted). Formal thresholds of statistical significance are added above each error bar to facilitate effect estimation (“+”: p<0.1; “*”: P<0.05; “**”: P<0.01; “***”: P<0.001).

Overall, country-level parameters produced effects of larger magnitude (Fig 2). Indeed, we observed sharp differences between countries with regard to the risk of duplication (Fig 3). Compared to the United States, the risk was significantly higher in China, India, Argentina and other developing countries (i.e. all those included in the “other” category, Fig 3). Multiple other countries (e.g. Belgium, Austria, Brazil, Israel, etc.) also appeared to have higher average risk than the United States but the very small number of studies from these countries hampered statistical power and thus our ability to draw any conclusion. Germany and Australia tended to have lower risk than the United States, but only Japan had a statistically significant lower risk (Fig 3).

**Fig 3:**
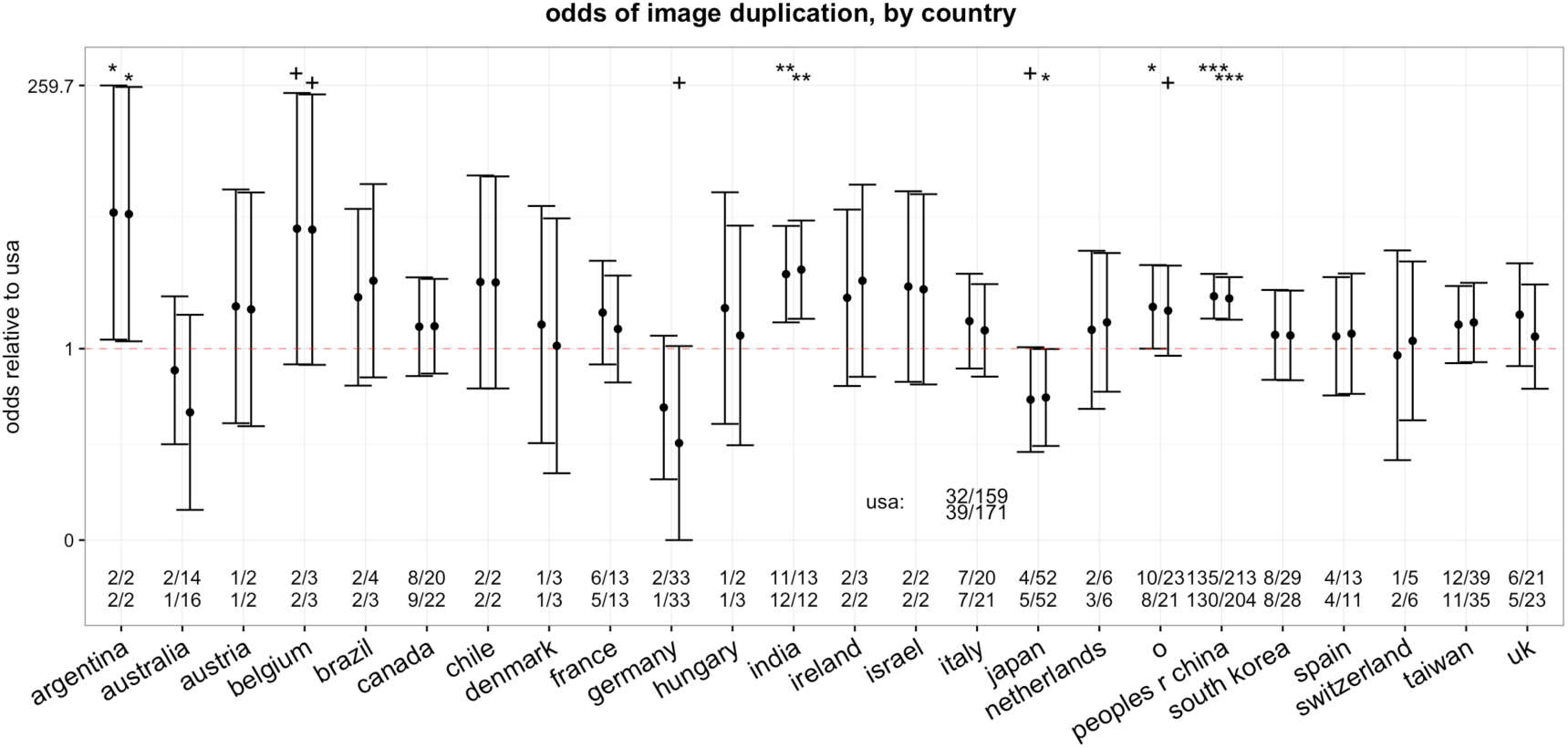
Effect (odds ratio and 95%CI) of country of activity of first and last authors on the likelihood to publish a paper containing a Category 2 or 3 problematic image duplication, compared to authors working the United States. The data were produced with a multivariable logistic regression model, in which dummy variables are attributed to countries that were associated with the first or last author of at least one treatment and one control paper. All other countries were included in the “other” category. Numeric data are raw numbers of treatment and control papers for first and last author (upper and lower row, respectively). Formal thresholds of statistical significance are added above each error bar to facilitate effect estimation (“+”: p<0.1; “*”: P<0.05; “**”: P<0.01; “***”: P<0.001).

To reduce the possible confounding effect of country, we performed secondary analyses on subsamples of countries with relatively homogeneous cultural and economic characteristics (Fig S1). Such sub-setting appeared to improve the detection of individual-level variables. In particular, the risk of duplication appeared to be positively associated with authors’ publication rate, citation score, journal score and female gender (Fig S1 a-h, and see SM for all numerical results). These effects, however, were never formally statistically significant in such univariable analyses.

Secondary multivariable analyses, however, corroborated all of our main results (Fig 4). A model that included individual parameters, as well as an interaction term between number of authors and number of countries (in place of the country-to-author ratio, which is not independent from the number of authors) and country-level parameters of publication and misconduct policies suggested that the risk of misconduct was predominantly predicted by country and team characteristics (Fig 4a). The risk was significantly higher in countries with cash-based publication incentives, lower in those with national misconduct policies, and grew with team size as well as with number of authors, with the latter two factors modulating each other: for a given distance, larger teams were less at risk from misconduct, as the social control hypothesis predicted (Fig 4a).

**Fig 4:**
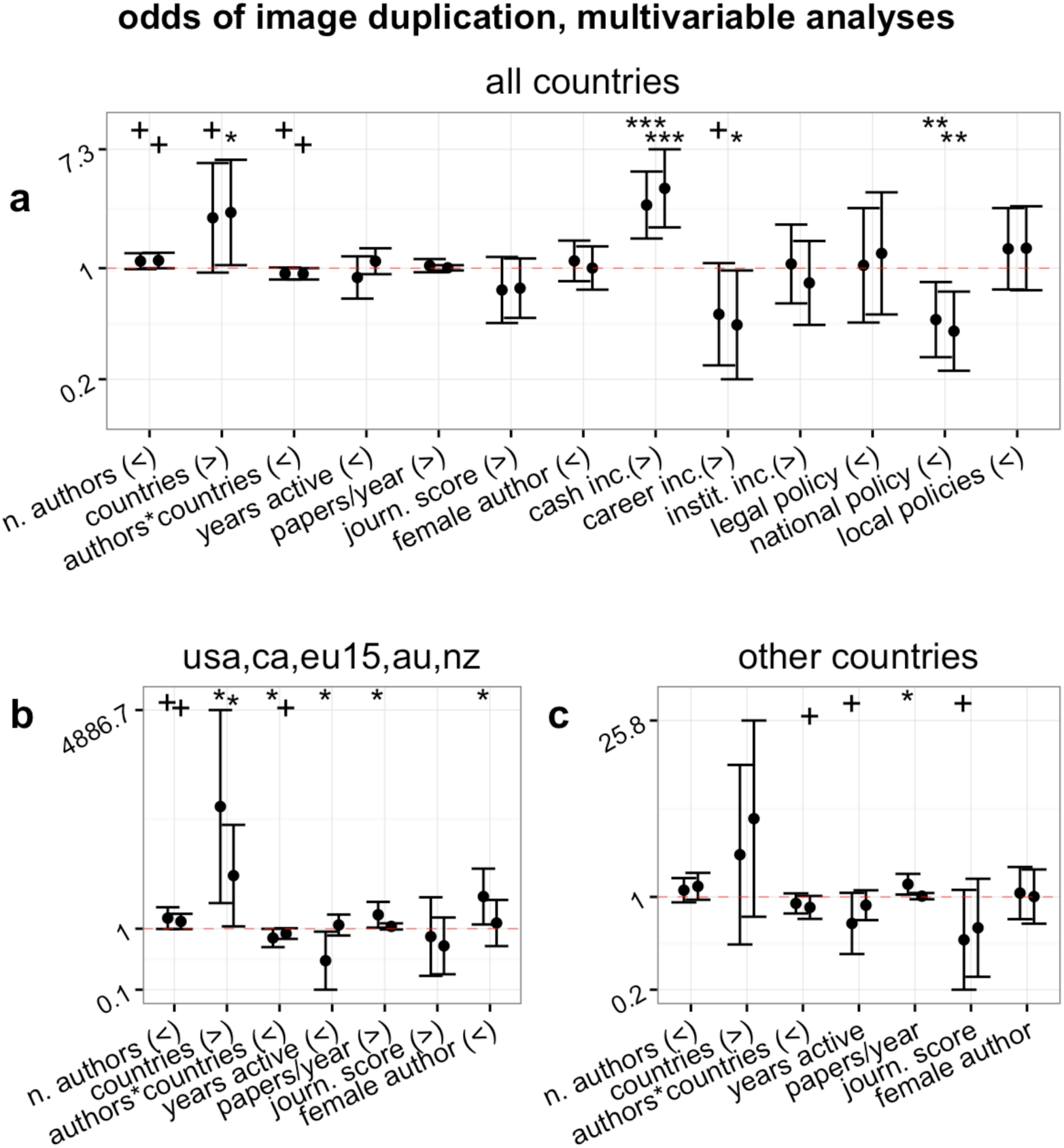
Effect (odds ratio and 95%CI) of characteristics of study and first and last author on the probability of publishing a paper containing a Category 2 or 3 problematic image duplication. Each subpanel illustrates results of a single multivariable model, partitioned by country subsets (see text for further details). First and second error bars correspond to data from first and last authors, respectively. Signs in parentheses indicate direction of expected effect (“>” : OR>1; “<” : OR<1). Formal thresholds of statistical significance are added above each error bar to facilitate effect estimation (“+”: p<0.1; “*”: P<0.05; “**”: P<0.01; “***”: P<0.001).

When limited to English-speaking and EU15 countries, multivariable analyses of individual and team characteristics supported most theoretical predictions, suggesting that misconduct was more likely in long-distance collaborations and amongst early-career, highly productive and high-impact first-authors (Fig 4b). Female first authors were significantly more at risk of being associated with Category 2 and 3 problems, a finding that is inconsistent with the gender hypothesis. Analyses on the remaining subset of countries yielded similar results (Fig 4c).

Almost identical results were obtained with a non-conditional logistic regression model, consistent with the fact that our sample was homogeneous with regards to important characteristics such as journal, methodology and year of publication. Results obtained combining all three categories of duplications were largely overlapping with those presented in the main text and would have led to similar conclusions (see all numerical results in SI).

## DISCUSSION

To the best of our knowledge, this is the first direct test of hypotheses about the causes of scientific misconduct that was conducted on an unbiased sample of papers containing flawed or fabricated data. Our sample represented papers containing honest errors and intentional fabrications of various degrees in unknown relative proportions. However, we correctly predicted that Category 1 duplications would exhibit smaller or null effects, whilst most significant effects, if observed at all, would be observed in Categories 2 and 3 (Fig 1). Support of this prediction retrospectively confirms that, as suggested by a previous analysis of these data [5], Category 1 duplications are most likely the result of unintentional errors or flawed methodologies, whilst Category 2 and 3 duplications are likely to contain a significant proportion of intentional fabrications.

Results obtained on Category 2 and 3 papers, corroborated by multiple secondary analyses (see SI), supported some predictions of the hypotheses tested, but did not support or openly contradicted others:

- *Pressure to publish hypothesis:* partially supported. Early-career researchers, and researchers working in countries where publications are rewarded with cash incentives were at higher risk of image duplication, as predicted. However, countries having other publication incentive policies had a null or even negative risk (Fig 1b). In further refutation of predictions, individual publication rate and impact of authors was not or negatively associated with image duplication, although in secondary multivariable analyses we observed a positive association between publication rate of first authors and risk of duplication. The latter finding might represent the first direct support of this prediction, but should be verified in future confirmatory tests. The correlation with cash incentives may not be taken to imply that such incentives were directly involved in the problematic image duplications, but simply that such incentives may reflect the value system in certain research communities that might incentivize questionable research practices.
- *Social control hypothesis:* supported. In univariable analyses, only predictions based on socio-cultural conditions of different countries were in large agreement with observations (Fig 1c). However, when country characteristics were controlled and/or adjusted for, we observed a consistent negative interaction between number of authors and number of countries per author in a paper, which is in good agreement with the hypothesis (Fig 4).
- *Misconduct policy hypothesis:* partially supported. Countries with national and legally enforceable policies against scientific misconduct were significantly less likely to produce image duplications (Fig 1e, Fig 4a). However, other misconduct policy categories were not associated with a reduced risk of image duplication, and tended if anything to have a higher risk. As noted above for publication incentive policies, we cannot prove a cause-effect relationship. The presence of national misconduct policies may simply reflect the greater attention that a country’s scientific community pays to research integrity.
- *Gender hypothesis:* not supported. In none of the main and secondary analyses did we observe the predicted higher risk for males. Some of the secondary analyses might have found an association between female authors and the risk of image duplication (Fig 4b). This latter finding, however, needs to be validated in future confirmatory studies.

A previous, analogous analysis conducted on retracted and corrected papers had led to largely similar conclusions [12]. The likelihood to correct papers for unintentional errors was not associated with most parameters, similarly to what this study observed for category 1 duplications. The likelihood to retract papers, instead, was also found to be significantly associated with misconduct policies, academic culture, as well as early-career status and average impact score of first or last author. Differently from what this study observed on image duplications, however, individual publication rate was negatively associated with the risk of retraction and positively with that of corrections [12]. We hypothesize that at least two factors may underlie this difference in results. First, analyses on retractions included every possible error and form of misconduct, including plagiarism, whereas the present analysis is dedicated to a very specific form of error or manipulation. Second, analyses on retractions are intrinsically biased and subject to many confounding factors, because retractions are the end results of a complex chain of events (e.g. a reader signals a possible problem to the journal, the journal editor contacts the author, the author’s institution starts an investigation, etc….) which can be subjected to many sources of noise and distortion. Therefore, whilst on the one hand our results may be less generalizable, on the other hand they are more accurate and less biased than results obtained on retractions.

A remarkable agreement was also observed between these results and those of a recent assessment of the causes of bias in science, authored by two of us [14]. This latter study tested similar hypotheses using identical independent variables on a completely different outcome (the likelihood to over-estimate results in meta-analysis) and using a completely different study design. Therefore, the convergence of results with this latter study is even more striking and strongly suggests that all these separate analyses are detecting genuine underlying patterns that reflect a connection between research integrity and characteristics of authors, team and country.

The present study has avoided many of the confounding factors that limit studies on retractions, but could not avoid other limitations. An overall limitation concerns the kind of image duplication analyzed in this study, which is only one of the many possible forms of data falsification and fabrication that may occur in the literature. This restriction limits in principle broad generalizations. However, as noted above, our results are in large agreement with previous analyses that encompassed all forms of bias and misconduct [12, 14], which suggests that our findings are consistent with general patterns linked to these phenomena.

Two other possible limitations of our study design made results very conservative. Firstly, we could not ascertain which of the duplications were actually due to scientific misconduct and which ones derived from honest error, systematic error or negligence. Secondly, our individual-level analyses focused on characteristics of the first and the last author, under the assumption that authors in these positions are most likely to be responsible for any flaws in a publication. However, we do not know who, amongst the co-authors of included studies, was actually behind the problematic duplication. Both these limitations ought to increase confidence in our results, because they are likely to have reduced the magnitude of measurable effect sizes. As our univariable analyses confirmed, image duplications that are due not to scientific misconduct but to unintentional error are unlikely to be associated with any factor (Fig 1). Similarly, if an image duplication was not caused by its study’s first or last author, then we simply would not expect the characteristics of first and last author to be associated with the likelihood of that error. Therefore, to any extent that they affected the study, these two limitations have introduced random noise in our data, reducing the magnitude of any measurable effect and thus making our results more conservative.

Any random noise in our data might have reduced the statistical power of our analyses, for the reasons discussed above. However, our statistical power was relatively large. Even when restricted to the smallest subset (e.g. category 3 duplications) our analyses had over 89% power to detect an effect of small magnitude. We can therefore conclude that, despite the limitations discussed above, all of our tests had sufficient power to reject null hypotheses for at least large and medium effect sizes.

A further possible limitation in our analysis pertains to the accuracy with which we could measure individual-level parameters. Our ability to correctly classify the gender and to reconstruct the publication profile of each author was subject to standard disambiguation errors [27] which may be higher for authors in certain subsets of countries. In particular, authors from South- and East-Asian countries have names that are difficult to classify, and often publish in local journals that are not indexed in the Web of Science and were therefore not captured by our algorithms. Any systematic bias or error in quantifying parameters for authors from these countries would significantly skew our results because country-level factors were found in this study - as well in previous studies on retractions - to have significant effects [12]. However, all our main conclusions are based on effects that were measured consistently in subsets of authors based on countries at lower risk of disambiguation error. Moreover, this limitation is only likely to affect the subset of tests that focused on author characteristics.

Indeed, this study suggests that significant individual-level effects might not be detectable unless country-level effects are removed or adjusted for. This prominence of country-level effects in determining the risk of problematic image duplications might be one of the most important finding of this study. We observed clear and indisputable evidence that problematic image duplications are overwhelmingly more likely to come from China, India and other developing countries, consistent with the original interpretation of these data [5]. Regardless of whether the image duplications that we have examined in this study were due to misconduct or unintentional error, country-level effects suggest that particular efforts might be needed to improve the reliability of studies from developing countries.

Previous analyses on retractions, corrections and bias [12, 14] as well as the present analysis of image duplications cannot demonstrate causality. However, all these analyses consistently suggest that developing national misconduct policies and fostering an academic culture of mutual criticism might be effective preventive measures to ensure the integrity of future research.

## MATERIALS AND METHODS

Methods of this study very closely followed the protocol of a previous analysis of risk factors for retractions and corrections [12]. To guarantee the confirmatory and unbiased nature of our analyses, all main and secondary analyses as well as sampling and analytical methodology were pre-specified and registered at the Center for Open Science (osf.io/w53yu) [28]. The main goal of the analysis was to produce a matched-control retrospective analysis aimed at identifying which characteristics of papers and their authors were significantly predictive of the likelihood to fall into the “treatment” as opposed to “control” category (papers with or without problematic image duplications, respectively).

### Sampling of papers

Papers had been identified by the independent assessment of three of the present paper’s authors (EB, AC, FF). Control papers were retrieved from the set of papers that had been examined by the authors and in which no evidence of data duplication had been recognized. For each treatment paper, two controls were retrieved for inclusion, i.e. one published immediately before and one immediately after the treatment paper. Order of publication was determined based on Web of Science’s unique identifier code. When the candidate control paper of one treatment paper coincided with the candidate control of another treatment paper, the next available control paper was selected instead.

### Data collection

Following previous protocols[12, 14], we collected a set of relevant characteristics of all included papers and of all of their authors. More specifically, for each paper we recorded:

- Number of authors of each paper.
- Number of countries listed in the authors’ addresses.
- Average distance between author addresses, expressed in thousands of kilometers. Geographic distance was calculated based on a geocoding of affiliations covered in the Web of Science [29].

For each author of each paper in the sample we retrieved the following data from the Web of Science:

- Year of first and last paper recorded in the Web of Science.
- Total number of article, letters and review papers authored or co-authored.
- Total number of citations received by all papers authored or co-authored.
- Field-normalized citation score.
- Field-normalized journal impact score.
- Proportion of papers authored or co-authored that appeared in the top-10 journals of that author’s field.
- Author’s main country of activity, based on the address most commonly indicated.
- Author’s first name. The combination of first name and country was used to assign gender. The majority of gender assignments were made by a commercial service (genderapi.com) but an attempt was made to identify the gender of unassigned names. When neither approach could attribute an author’s gender reliably, gender was assigned to the “unknown” category.

Country information was used to assign each author to the corresponding country-level variable, using the following scheme:

- Publication incentives policies: i.e. cash-incentives to individuals (CN, KR, TU); performance linked to individual’s career (DE, ES, USA); performance linked to institution’s funding (AU, BE, NZ, DK, IT, NO, UK), based on classifications in [30].
- Social control hypothesis: developmental state - German academic model (CN, JP, KR); intermediate case (DE, SI, TW, ISR); regulatory state & Anglo-American academic culture (US, UK), based on classification by [16].
- Misconduct policy: national and legally enforced (USA, DK, NO); national non-legally enforced (UK, SW, FI, NL, DE, AT, AU, JP, CN, KR, CR, TN, ZA); local (institutional) policies (ES, IL, FR, BE, CH, EE, LV, PL, CZ, HU, PE, GR, IN, BD), data based on references in [12].

Although we collected information for all authors of the papers, we only tested individual predictors measured on the first and last authors, positions that in biomedical papers tend to be attributed to the authors that most contributed to the research, often in the role of junior and senior author, respectively [31, 32].

### Analyses

All variables were included in the analysis untransformed, although a few variables were re-scaled linearly: author publication rate data was divided by 10, geographic distance data was divided by 1000, and countries-to-author ratio was multiplied by 100. This re-scaling of some variables served the purpose of improving the visibility of effect sizes in figures and had no impact on the results.

All hypotheses were tested using standard conditional logistic regression analysis, i.e. a logistic regression model with an added “stratum” term that identifies each subgroup of treatment and matched controls. The conditional logistic regression approach is most useful when papers differ widely in important characteristics, such as year and journal of publication (see [12]). Analyses were also repeated with a non-conditional logistic regression to assess the robustness of the results. Analyses were conducted with all three categories of duplication combined, separately on category 1 and category 2 and 3 papers, and combining categories 2 and 3.

Since the sample size was pre-determined, we did not conduct a prospective power analysis. Post-hoc power analyses based on unconditional logistic regression suggest that our main analyses, when combining papers from all duplication categories (a total of 1039 data points) had over 99% statistical power to detect a small effect size (i.e. OR=1.5), and 89% power for analyses restricted to the smallest subsample, i.e. category 3 duplications. All analyses were conducted with the open-source statistical package Survival implemented by the R software [33].

## ACKNOWLEDGEMENTS. AUTHOR CONTRIBUTIONS

Conceived and designed study: DF; Collected data: RC, EB; contributed reagents and materials: EB, FF, AC; analyzed data: DF; wrote the paper: DF with input from all authors.

---

**Fig S1:**
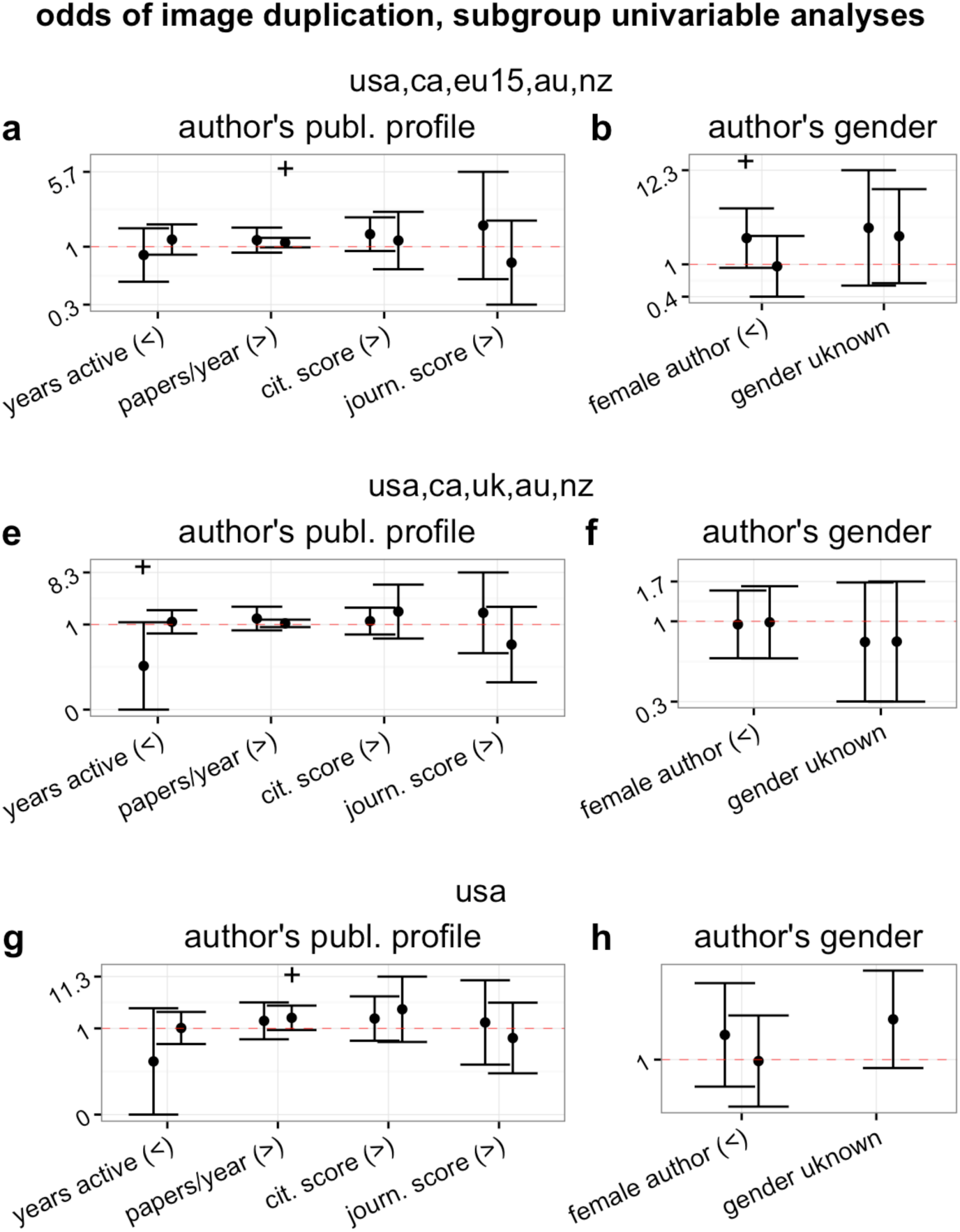
Effect (odds ratio and 95%CI) of characteristics of first and last author on probability of publishing a paper containing a Category 2 or 3 image duplication. Each subpanel shows results of univariable analyses on subsets of countries (see text for further details). First and second error bar correspond to data from first and last authors, respectively. Panels are subdivided according to overall hypothesis tested, and signs in parentheses indicate direction of expected effect (“>” : OR>1; “<” : OR<1). Formal thresholds of statistical significance are added above each error bar to facilitate effect estimation (“+”: p<0.1; “*”: P<0.05; “**”: P<0.01; “***”: P<0.001).

